# Interspecific Aggression Of Two Wrens Along An Environmental Gradient In Western Ecuador

**DOI:** 10.1101/2023.10.13.562238

**Authors:** Luis Daniel Montalvo, Rebecca T. Kimball, Scott K. Robinson

## Abstract

Interspecific territoriality is a prevalent form of interference competition among animals. However, the connections between hybridization, climate, and interspecies territorial aggression in tropical regions remain largely unexplored. Here, we investigated territorial aggression in two hybridizing tropical bird species, *C. z. brevirostris* and *C. f. pallescens*, in western Ecuador using playback experiments. We tested three hypotheses: 1) hybridizing species exhibit comparable intra- and inter-specific territorial aggression; 2) asymmetrical aggression driven by *C. z. brevirostris* dominance determines gene flow patterns; and 3) precipitation influences territorial aggression. Supporting hypothesis 1, the admixed *C. f. pallescens* North showed no difference in intra-vs inter-specific aggression. However, the non-admixed *C. f. pallescens* South exhibited greater inter-specific aggression, providing partial support for hypothesis 1. Contrary to hypothesis 2, *C. f. pallescens* South displayed significantly higher aggression than *C. z. brevirostris* and *C. f. pallescens North*. Furthermore, precipitation models outperformed null models, supporting hypothesis 3 that precipitation influences *Campylorhynchus* territorial aggression. Collectively, these findings suggest hybridization can stabilize coexistence via territoriality, and precipitation strongly affects aggression, potentially through resource availability. Unexpectedly, *C. z. brevirostris* dominance did not appear to drive asymmetric introgression between species, warranting further investigation into the underlying mechanisms. Complex factors shape territorial aggression in tropical birds, including genetic admixture, group size, latitude, and climate. This study highlights the need for additional research elucidating the relationships between hybridization, territoriality, and environmental stressors in tropical avian communities. We discuss possible mechanisms explaining the detected effects of precipitation on aggression and the lack of *C. z. brevirostris* dominance in determining introgression patterns.

## INTRODUCTION

Interspecific territoriality is one of the most prevalent types of interference competition in animals (Simmons 1951, Cody 1973, Peiman and Robinson 2010, Grether et al. 2017), and has been found to modify distribution ranges, especially where there is strong asymmetric aggression between species at borders. (Case et al. 2005, Price and Kirkpatrick 2009, Jankowski et al. 2010, Pasch et al. 2013, Taylor et al. 2015). Two main foundational hypotheses concerning the origin and continuity of interspecific territoriality posit that it could emerge as a by-product of intraspecific territoriality when allopatric species undergo secondary contact and maintain similar signaling, in which case interspecific territoriality is considered “mal-adaptive” (Cowen et al. 2020, Drury et al. 2020). The second hypothesis posits its evolution between species for the same reasons as intraspecific territoriality: interspecific territoriality in birds usually persists because of interspecific competition for mates, resources, or both (Cowen et al. 2020, Drury et al. 2020).

In the latter hypothesis, interspecific territoriality may stabilize the local coexistence of closely related and strong resource competitors or hybridizing species, while enabling a faster transition into sympatry than without it (Cowen et al. 2020, Drury et al. 2020). For instance, two allopatric species with extensive resource overlap could coexist in sympatry without diverging in resource use by engaging in interspecific territoriality, thereby spatially partitioning the resources (Cowen et al. 2020). Hybridizing species are more likely to be interspecifically territorial than non-hybridizing ones, making hybridization a crucial predictor of interspecific territoriality even for species pairs with relatively low breeding habitat overlap (Drury et al. 2015, 2020). Understanding which of these hypotheses explains interspecific territoriality is crucial for developing a general comprehension of how it affects the local coexistence and evolutionary dynamics of interacting species (Cowen et al. 2020).

Here, we explore the hypothesis that hybridizing species occurring in syntopy, and similar niches tend to exhibit interspecific territoriality. To accomplish this, we measured intra- and interspecific territorial aggressive responses of *C. z. brevirostris*, *C. fasciatus*, and genetically admixed populations along a precipitation gradient in Western Ecuador. The genus *Campylorhynchus* is known for its group-territorial aggressive behavior (Anderson and Anderson 1959, Lindell 1996, Bradley and Mennill 2009). There is evidence that populations of *C. fasciatus* in Southwest Ecuador and Northwest Peru (*C. f. pallescens* South) are genetically distinct from an admixed populations at the environmental transition in western Ecuador (*C. f. pallescens* North) (Montalvo et al. in prep) and there is potential higher gene flow from *C. z. brevirostris* towards *C. f. pallescens* North (Montalvo et al. in prep).

In our first hypothesis (H1), we tested if the population with higher admixture (*C. f. pallescens* North) exhibits similar levels of intra- and interspecific territorial aggression while more genetically differentiated populations (*C. z. brevirostris* and *C. f. pallescens* South) show significantly more intraspecific aggression. In our second hypothesis (H2), we tested if aggressive territoriality plays a role in driving gene flow, if this is true then we should expect that *C. z. brevirostris* would be dominant over *C. f. pallescens* North and South. Hybrid zones are regions where genetically distinct populations meet and produce offspring and can be found across environmental gradients (Harrison and Larson 2014), such as the one along Western Ecuador. These zones are particularly useful systems for assessing the role of behavior in maintaining the ranges of overlapping, interbreeding taxa (Harrison 1993).

Climate variability has a significant impact on the dynamics of hybrid zones (Gee 2004, Taylor et al. 2015, Guillaumet and Russell 2022) and interspecific territorial aggression (Jankowski et al. 2010, Bishop et al. 2015, Freeman et al. 2017). Interspecific aggression and competition can constrain elevational ranges, and climate changes can lead to range shifts, species loss, and altered ecosystem functions provided by birds (Jankowski et al. 2010, Charmantier and Gienapp 2014). Although the specific mechanisms through which climate variability impacts birds’ behaviors and distributions are not well understood, climate variability has been shown to affect habitat quality (Barnagaud et al. 2012, Jackson et al. 2015), which can affect territoriality in birds (Robinson and Terborgh 1995, Ekman et al. 2001, Carrete et al. 2006, Jankowski et al. 2010). Rainfall patterns, for example, can directly affect plant productivity and insect abundance, which in turn determine food availability for birds (Haddad et al. 2001, Zhu et al. 2014, Smith and Williams 2023). The quality of bird habitats in low-precipitation regions is influenced by a variety of factors, including food availability, substrate diversity, and morphological adaptations to specific habitats (Smith et al. 2010, Akresh et al. 2019, Tesfahun and Ejigu 2022). It has been suggested that in high-quality habitats, where food resources are abundant, competition for food is reduced, resulting in a lower intensity of competition (Dhondt 2010, Scales et al. 2011). In this case, more dominant species are expected to occur in the most productive habitat (Robinson and Terborgh 1995, Jankowski et al. 2010). If these findings are generalizable to Campylorhynchus along Western Ecuador, then *C. z. brevirostris*, which occurs in higher-quality habitats (wet region), should exhibit higher levels of territorial aggression and dominance, which is consistent with hypothesis H2.

Additionally, we explored the association of territorial aggression in *Campylorhynchus* and spatially explicit climate variability along the precipitation gradient in Western Ecuador. We hypothesized (H3) that if the climate has a direct or indirect effect on habitat quality, and habitat quality influences aggressive territoriality, then the aggressive territoriality of *Campylorhynchus* should correlate with spatial variation of climate along Western Ecuador. Studying underlying promotors and modulators of territorial aggression allows us to understand the coping mechanisms of biodiversity to climate change (Jankowski et al. 2010, Cockrem 2013, Elsen et al. 2017, Grether and Okamoto 2022).

The present study aims to address the knowledge gap concerning the link between hybridization, climate, and interspecies territorial aggression in tropical regions – a relationship that remains largely unexplored in the field — and contributes to our understanding of the factors that contribute to species diversity and evolution.

## METHODS

### Study Area

We carried out playback experiments in 12 sampling sites along Western Ecuador (Figure 1). We considered breeding groups to belong to a different sampling location if they were at least 20 km apart from the closest location. Western lowland Ecuador is characterized by a moisture gradient that goes from the humid Choco-Darien-Western Ecuador region in the north to the dry Tumbesian region in the south. Rainfall in the northern part ranges from 2000 to 7000 mm annually, while the southern part of the region, with less than 1000 mm annually, has an eight-month dry season (Dodson and Gentry 1991).

**Figure 1.**
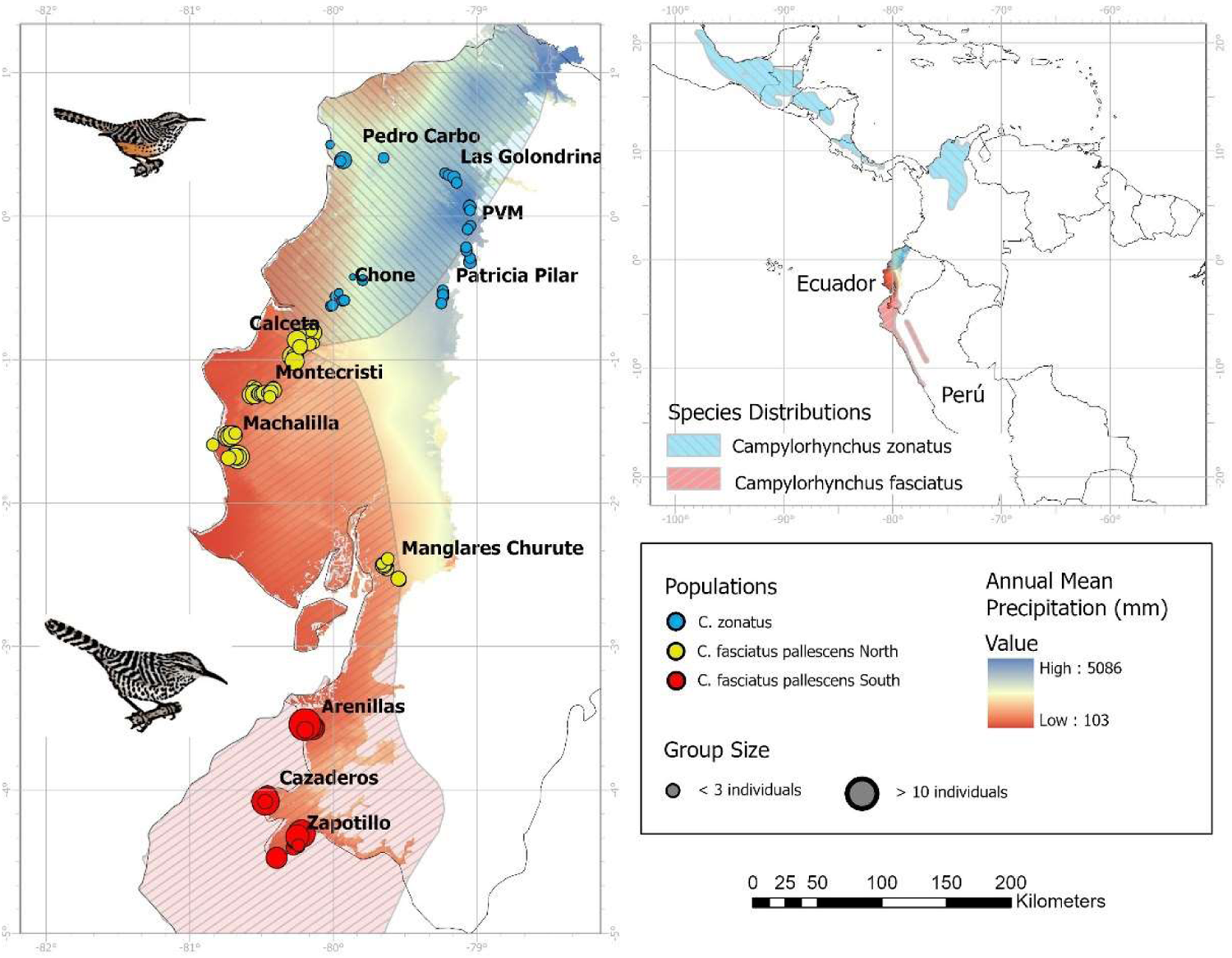
Groups and sampling sites along the precipitation gradient in western Ecuador. Genetically differentiated populations (Montalvo et al. in review) are represented with circles of different colors. Size of circles represent number of individuals in groups. Species distributions according to Ridgely and Greenfield (2006).

### Species System

Phylogenetic relationships of *Campylorhynchus* now available show that *C. fasciatus* and *C. z. brevirostris* are closely related but not sister species (Barker 2007, Burleigh et al. 2015, Vázquez-Miranda and Barker 2021). Both species have parapatric distributions along the precipitation gradient in west Ecuador, where *C. z. brevirostris*, occurring in lower abundance, is restricted to the wet region and *C. fasciatus* to the dry region (Ridgely and Greenfield 2001) (Figure 1). Both species feed mostly on insects and forage in dense vegetation on edges and clearings (Ridgely and Greenfield 2006, Kroodsma and Brewer 2020a, 2020b), which imply some degree of niche overlap.

### Playback Experiments

We designed and tested the experiments in 2017 during 39 preliminary playbacks in which we familiarized ourselves with the behavioral repertory and defined the response variables. It is known that *C. zonatus* and *C. fasciatus* are cooperative breeders and are generally observed in groups or related individuals (Kroodsma and Brewer 2020a, 2020b). Behaviors detected during the preliminary playbacks were performed by both adults with different degrees of juvenile participation. This territorial behavioral pattern allowed us to use the breeding group as a sample unit contrary to usual playback experiments where single individuals are used. We defined and classified fixed action patterns directed toward the source of stimuli (FAP) and constructed a catalog of territorial aggressive behaviors (Table 1). During these preliminary playbacks, we also recorded songs from the most northern and southern sampling locations possible (Table 2) for *C. zonatus* and *C. f. pallescens* South respectively representing parental populations. We selected eight songs for *C. zonatus* and nine for *C. fasciatus*. Except for eight experiments when we utilized recordings with three individuals, all recordings used in the experiments contained two birds singing simultaneously to control for this factor. We selected randomly what recordings to play during the experiments. We didn’t digitally modify the recordings except for volume and cutting vocalizations of other species when they did not overlap with the songs of *Campylorhynchus*. We calibrated the volume of recordings at 80 dB measured at one meter from the speaker using a BAFX3370 decibel meter.

**Table 1.** Category, code, and observed frequency of Fixed Action Patterns (FAP) directed to the speaker for interspecific territorial aggressive behaviors of *C. z. brevirostris* and *C. fasciatus*.

| Category | FAP | Code | Description | Frequency |
| --- | --- | --- | --- | --- |
| Sound | Bill clapping | BC | Produces noise by clapping the bill, normally lasts less than one second in duration | Medium |
| Sound | Calling | Cl | Produces short and simple sounds, at least two types were recognized, a rasp (tjak) and a ki sound | High |
| Sound | Singing | Si | Produces long and complex sound, it may last several seconds in duration and can be produced in duets | High |
| Movement | Hanging | Ha | Hangs with the head down | Low |
| Movement | Move the same tree | MT | Changing perch in the same tree involves hopping or short flights less than 1 m in distance within the same tree | High |
| Movement | Switching trees | ST | Changing perch moving to a different tree or making movements longer than 1 m in distance within the same tree, involves just flights | High |
| Movement | Switching trees and singing | STS | Involves the same action as "Switching Trees" but sings at the same time | High |
| Movement | Sally-strike | Sal | Makes a wing-powered, flycatcher-type motion after which the bird leaves and returns to the same perch, used as a direct attack to the speaker | Low |
| Movement | Stepping on speaker | Sk | Perches on the speaker, unusual | Low |
| Movement | Hooping | Ho | Small jumps n the ground, generally toward the speaker |  |
| Direct aggression | A direct attack on conspecific | DA | Makes a bill or claw hit at another bird with a jump or short flight, directed at an individual of the same group or another group that also responded to the playback, often occurs with a rasp calling | Low |
| No aggressive | Foraging | Fo | Leg-powered movement, walking along branches while reaching out for food items among vegetation | Medium |
| No aggressive | Grooming | Gr | Fixes feathers with the bill on itself or a conspecific | Low |
| No aggressive | Perching | Pe | No wing or leg-powered movement stays on the same perch | Low |
| No aggressive | Bill cleaning | BCl | Rubs the bill against a surface, usually after eating a food item | Low |
| No aggressive | Excretion | Po | Ejects feces while perching. This is not considered aggressive as it is also performed outside of a territorial aggressive context. | Low |

**Table 2.**
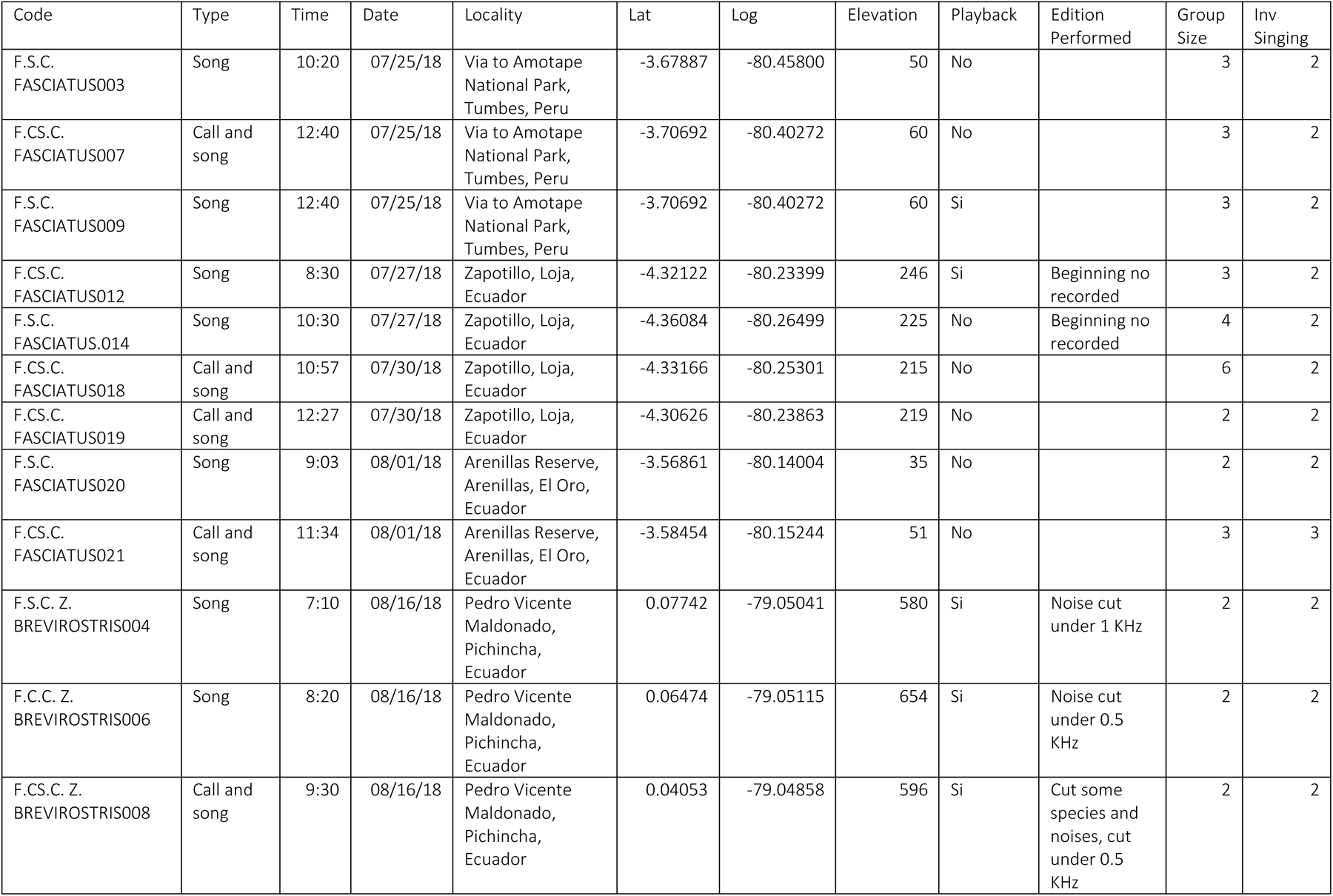
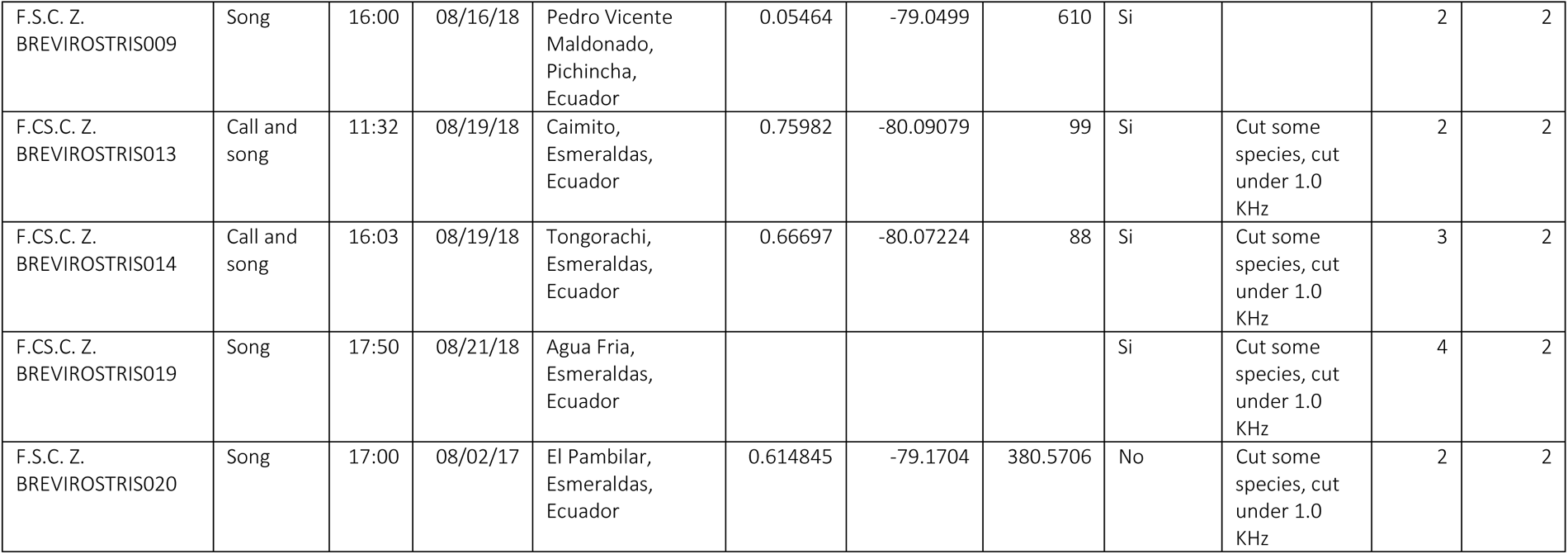
Metadata of vocalizations used in the playback experiments including code, types of vocalizations, time, date, locality (site, province, country), coordinates, if playback was used before the recording, digital editing done to the recordings, group size and the number of individuals singing in the recordings.

The data used for the analyses come from 112 playback experiments carried out between June and November of 2018. Breeding groups (a.k.a. groups) were tested only if they were at least 400 m apart from any other group already tested to maintain independence among group responses. Playback experiments were carried out after confirming auditorily or visually the presence of a group or individual. Recordings were played using a wireless UE BOOM 2 speaker. The observer kept a distance of at least 15 meters away from the speaker at the moment of the experiments. The experiments consisted of the following steps.

1. Initial observations. We annotated the context setting of the experiment including group size, the distance at which the group was first observed, other species present at the moment, habitat description, and coordinates.
2. Control. We played three minutes of the songs of *Elaenia flavogaster* (Yellow-bellied Elaenia) obtained from Xenocanto. *Elaenia flavogaster* occurs on edges and clearings along Western Ecuador (Ridgely and Greenfield 2006) in syntopy with both species.
3. Break 1. We recorded any change in the context setting different from the initial observation.
4. Treatment 1. We played three minutes of *C. z. brevirostris* or *C. f. pallescens* South alternating the order for each group encountered.
5. Break 3. We recorded changes in the setting or behaviors of the group.
6. Treatment 2. We played three minutes of *C. z. brevirostris* or *C. f. pallescens* South alternating the order according to what was played in treatment one.
7. Break 3. We registered the behaviors of the group if any were observed.

### Response Variables

For each group, we recorded all the FAPs (Table 1) observed during the experiment and the distance and time at which each FAP first occurred. We selected the closest distance to the speaker (MinDist) that the group, or any individual within it, achieved, the time since the beginning of the experiment to reach this distance (Latency), and the total number of FAPs observed per experiment and treatment (TFAP). We did not consider FAPs that were also observed frequently outside of experiments under not aggressive contexts (no aggressive in Table 1). We carried out a Principal Component Analysis (PCA) to summarize the variance of these three variables and used the first component (pca1) as a proxy of general territorial aggression and explored the association between response variables and pca1 using Pearson correlations and t-test to examine the significance of coefficients. Aggressive behaviors — as used in this study — are defined as actions that cause, threaten to cause, or seek to reduce physical damage, including threats, aggression, and submission (McGlone 1986).

### Explanatory Variables

We assigned each sampling site to one of the three genetically differentiated clusters (GCs) of *Campylorhynchus* along western Ecuador (Montalvo et al., in prep.) (Figure 1). In the high-quality habitat in the wet region and from North to South, sampling sites from Pedro Carbo to Chone were assigned to the parental GC of *C. z. brevirostris*, sampling sites from Calceta to Manglares Churute were assigned to the admixed GC of *C. f. pallescens* North, and sampling sites from Arenillas to Zapotillo were assigned to the GC of *C. f. pallescens* South (Figure 1).

We used climate variables from CHELSA 1.2 (Karger et al. 2017). CHELSA is a high-resolution (30 arc sec, ∼ 1km) free global climate data set. A multiple correlation analysis was used to find redundancies among these variables using Hmisc package for R (Harrell 2014). We selected climatic variables that we considered biologically relevant and showed low correlation coefficients with other selected variables to avoid collinearity. After this process, we selected Annual Mean Temperature (AMT), Annual Mean Precipitation (AMP), and Precipitation Seasonality (PS).

### Statistical Analysis

We carried out Bayesian Generalized Linear Models (BGLM) and a sampler-based Hamiltonian Markov Chain Monte Carlo (MCMC) available in the R package BRMS (Bürkner 2018). We tested hypotheses H1 and H2 by fitting a model (M1), for each response variable (MinDist, Latency, TFAP, pca1), that included fixed effects and interactions for treatment and genetic clusters. Hypothesis H1 predicts that there will be no significant differences between inter- and intraspecific treatments in groups belonging to the admixed genetic cluster of *C. f. pallescens* North, while there will be differences between intra- and interspecific treatments in *C*. *z. brevirostris* and *C. f. pallescens* South. Hypothesis H2 predicts that *C. z. brevirostris* will show significantly higher territorial aggression than the genetic clusters *C. f. pallescens* North and/or South. A posterior estimate was considered significant if the 95% confidence intervals (CI) did not contain zero.

To test hypothesis H3, for each response variable (MinDist, Latency, TFAP, pca1), we compared a null model with an intercept as a fixed effect (M0) to models that included the combined effect of all predictors (M7), as well as six models that included one of the following fixed effects: treatments and genetic clusters interactions (M1), group size (M2), latitude (M3), annual mean temperature (M4), annual mean precipitation (M5), and precipitation seasonality (M6). We carried out model selection to identify climate variables associated with the territorial aggression of *Campylorhynchus*. Our hypothesis H3 predicts that models including climate variables would be significantly better than the null model. We selected the best models by comparing the expected log pointwise predictive density (elpd) (Vehtari et al. 2017). If the elpd difference among models (elpd_diff) is less than four, then the models have very similar predictive performance (Sivula et al. 2022). If elpd_diff is larger than four, then compare that difference to the standard error of elpd_diff (Sivula et al. 2022). We obtained estimates of elpd using Bayesian leave-one-out cross-validation (LOO_CV) and Pareto smoothed importance sampling (PSIS) (Vehtari et al. 2017). The Pareto-k diagnostic (k̂) estimates how far an individual leave-one-out distribution is from the full distribution. If leaving out an observation changes the posterior too much, then importance-sampling is not able to give a reliable estimate. If k̂<0.5, then the corresponding component of elpd_loo is estimated with high accuracy. If 0.5<k̂<0.7 the accuracy is lower but still good. If k̂>0.7, then importance sampling is not able to provide a useful estimate for that component/observation (Vehtari et al. 2017). If there are many high k̂ values, we can gain more information from the effective number of parameters (p_loo). It is not needed for elpd_loo estimation but has diagnostic value. It describes how much more difficult it is to predict future data than observed data. In well-behaving cases p_loo<N and p_loo<p, where N is the sample size and p is the total number of parameters in the model (Vehtari et al. 2017). The model construction and model selection described above allow us to 1) test our hypotheses while reducing complexity, computational power, and the number of models to be tested, and 2) reduce overfitting when comparing a large number of models using LOO_CV. Overfitting could be an issue if many candidate models are being considered (Piironen and Vehtari 2017). We used default, weakly informative priors (improper flat priors for population-level effects and half-student t priors for group-level effects) for all models. The models ran for 100000 iterations, with a burn-in of the first 20000 samples. We compared visually the posterior predictive distributions against density distribution of observed data; if a model is a good fit, then we should be able to use it to generate data that looks a lot like the data we observed. The mixing and convergence of MCMC chains for all parameters were evaluated with the potential scale reduction factor (Ȓ) and visual inspections of MCMC diagnostic plots (Gelman and Rubin, 1992). The models for MinDist, and Latency were developed using an exponential distribution, TFAP was modeled using a Poisson distribution, while a Gaussian distribution was used for pca1, after testing for normality using Shapiro–Wilk test and exploring visually the density plots. All variables were scaled. We carried out all the analyses in R (R Core Team and R Development Core Team 2022).

## RESULTS

### PCA and Correlations among Response Variables

The PCA analysis summarizes 50% of the variation of the response variables in the first component (pca1) and 35% in the second component (pca2) (Figure 2). The pca1 is strongly correlated with MinDist (r=0.77, p-value<0.0001) and TFAP (r=-0.90, p-value<0.0001) while pca2 with Latency (r=-0.30, p-value< 0.0001). MinDist and TFAP were also significantly correlated (r=-0.51, pvalue<0.0001) (Figure 3).

**Figure 2.**
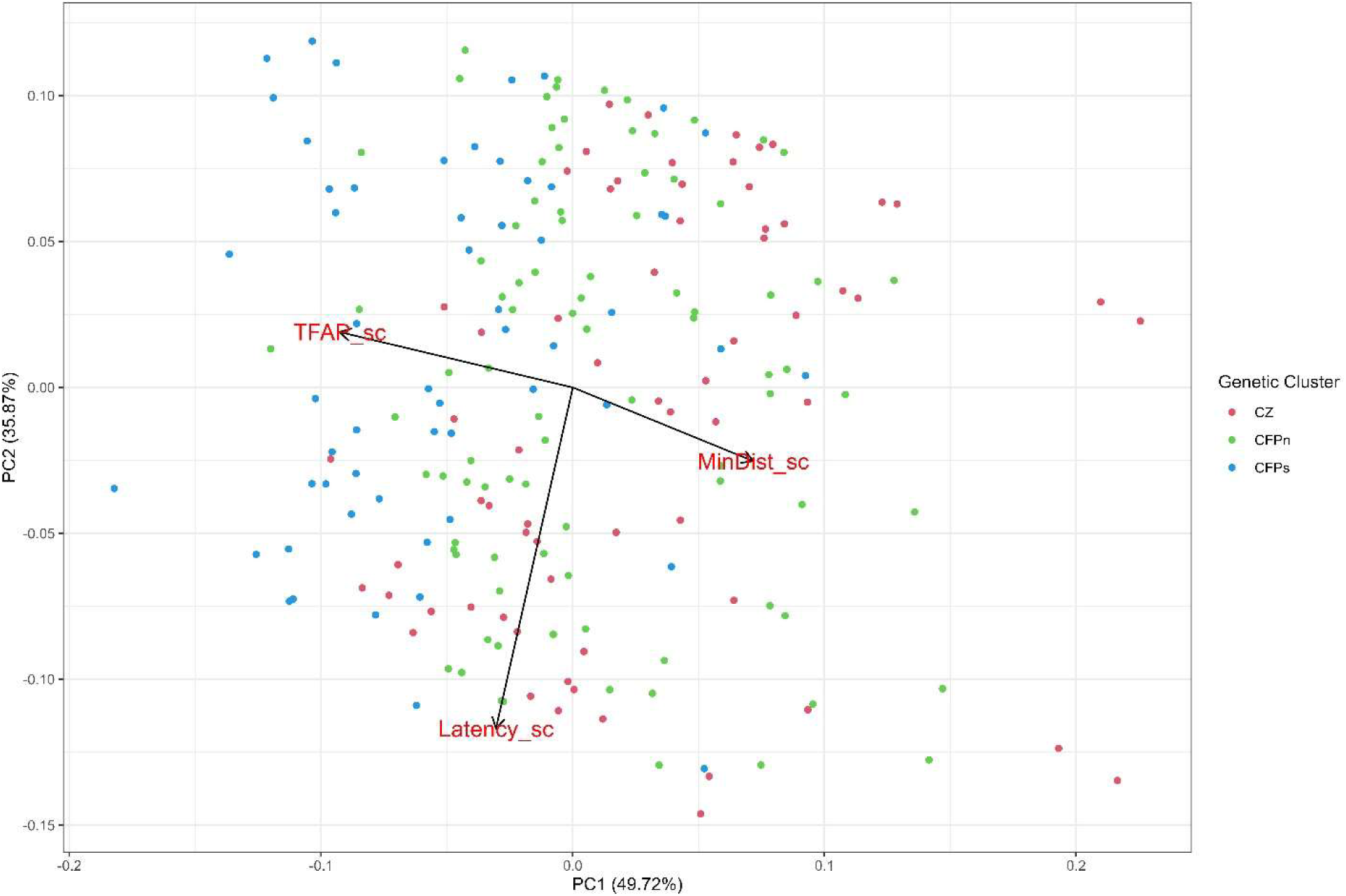
PCA of three aggressive response variables. MinDist_sc = scaled minimum distance reached by the group, Latency_sc = scaled time passed from the beginning of the experiment to reach minimum distance, and TFAP_sc = scaled total number of fixed actions patterns directed to the speaker. *C. f. pallescens* North is the population with higher admixture.

**Figure 3.**
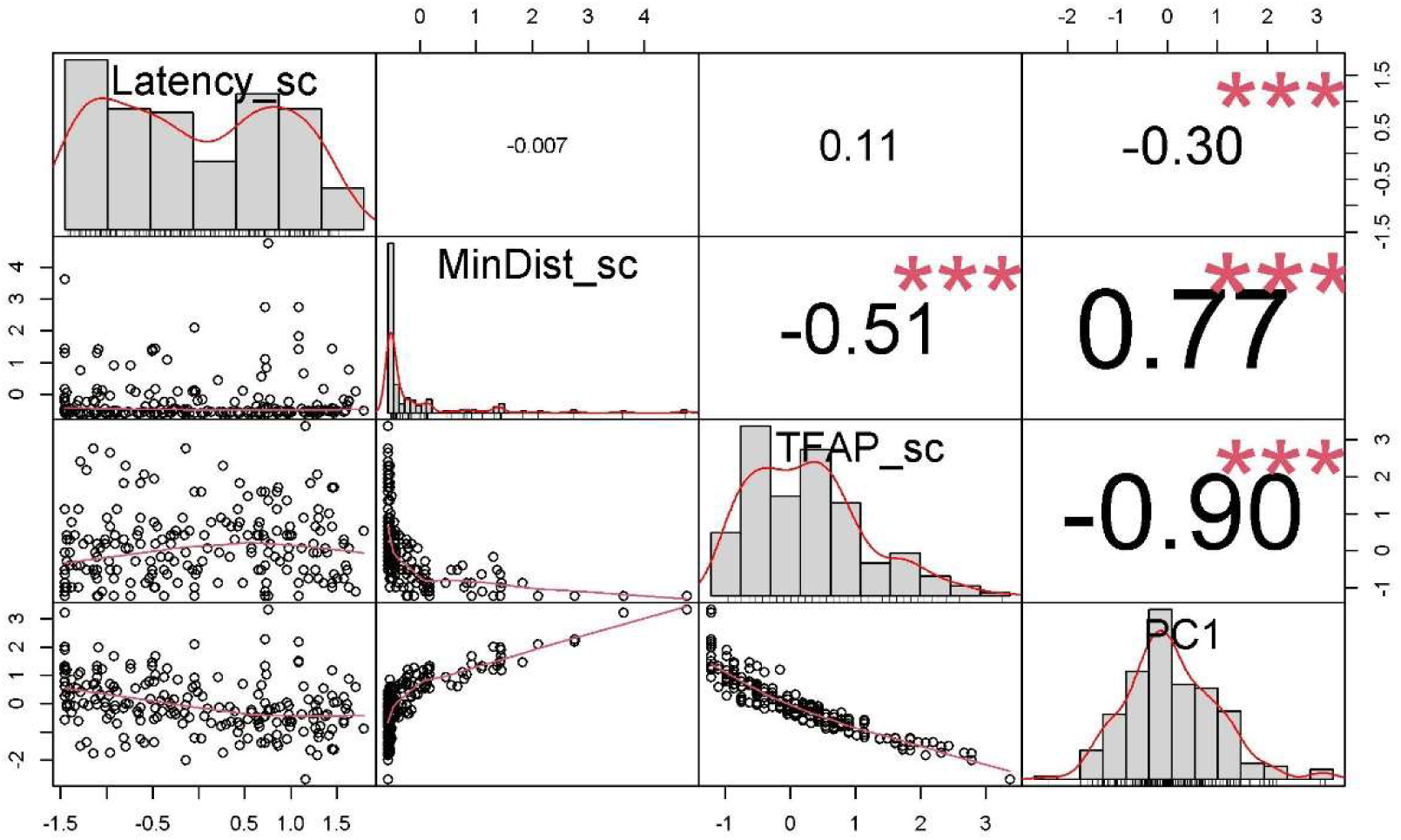
Pearson correlations among response variables. Each plot over the diagonal includes the correlation coefficients at the center while stars (***) represent p-values under 0.0001. Distributions of variables are shown on the diagonal. Variables plotted against each other are shown under the diagonal.

### Intra and Interspecific Territorial Aggression

No significant differences were observed between treatments and control for either of the genetic clusters concerning latency. Therefore, we will only refer to the results for minimum distance, TFAP, and pca1 in the remainder of this study.

Our analyses revealed no significant differences between intra- and interspecific treatments within the admixed genetic cluster *C. f. pallescens* North, which aligns with our initial hypothesis H1 (Figures 4-6). The 95% confidence intervals for both treatments included zero in this genetic cluster for all response variables examined (Minimum distance=0.34, 95% CI -0.31 to 1.02; TFAP=-0.04, 95% CI -0.26 to 0.17; pca1=0.20, 95% CI -0.45 to 0.84.

**Figure 4.**
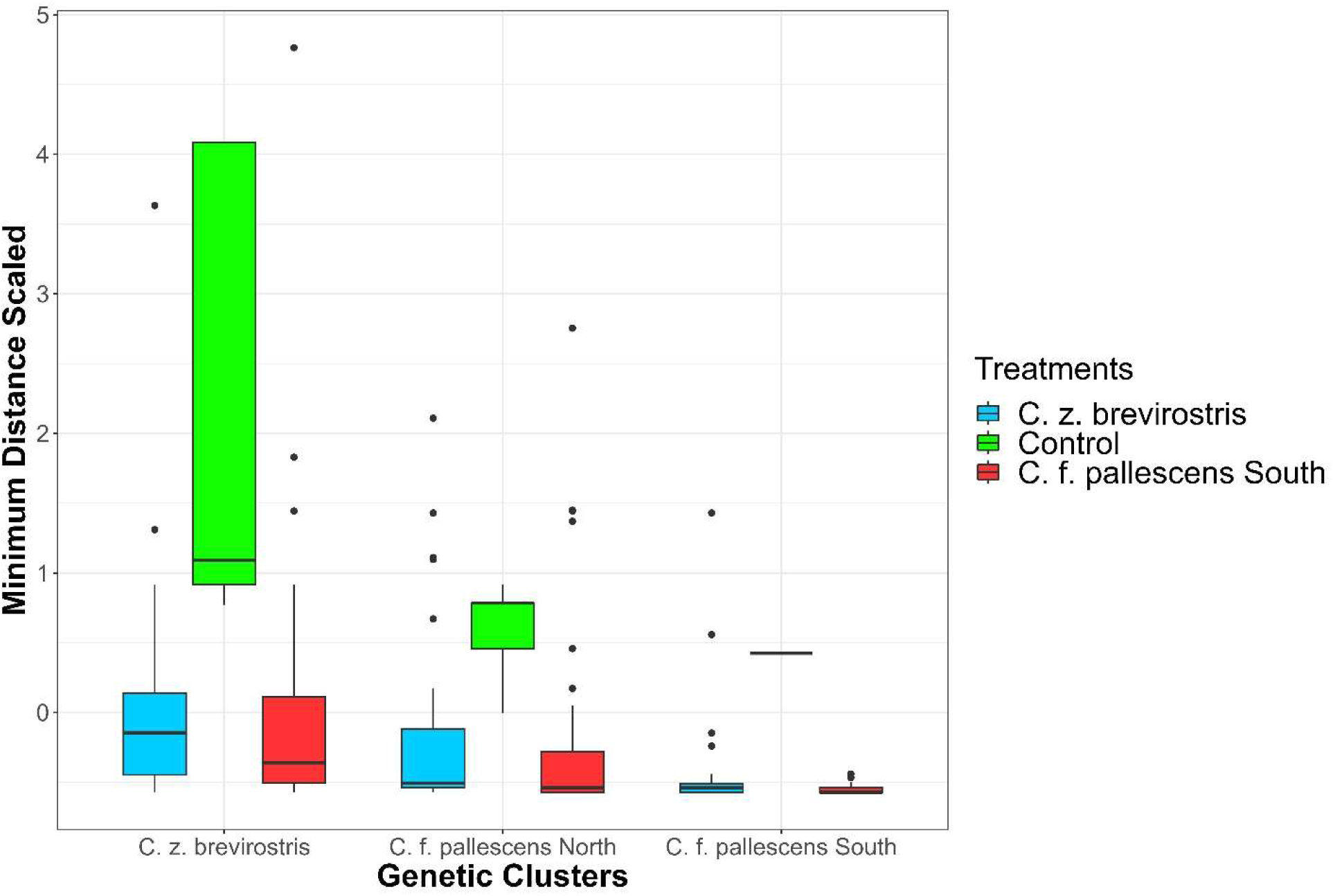
Median and 95% confidence intervals of minimum distance per treatment and genetic cluster.

**Figure 5.**
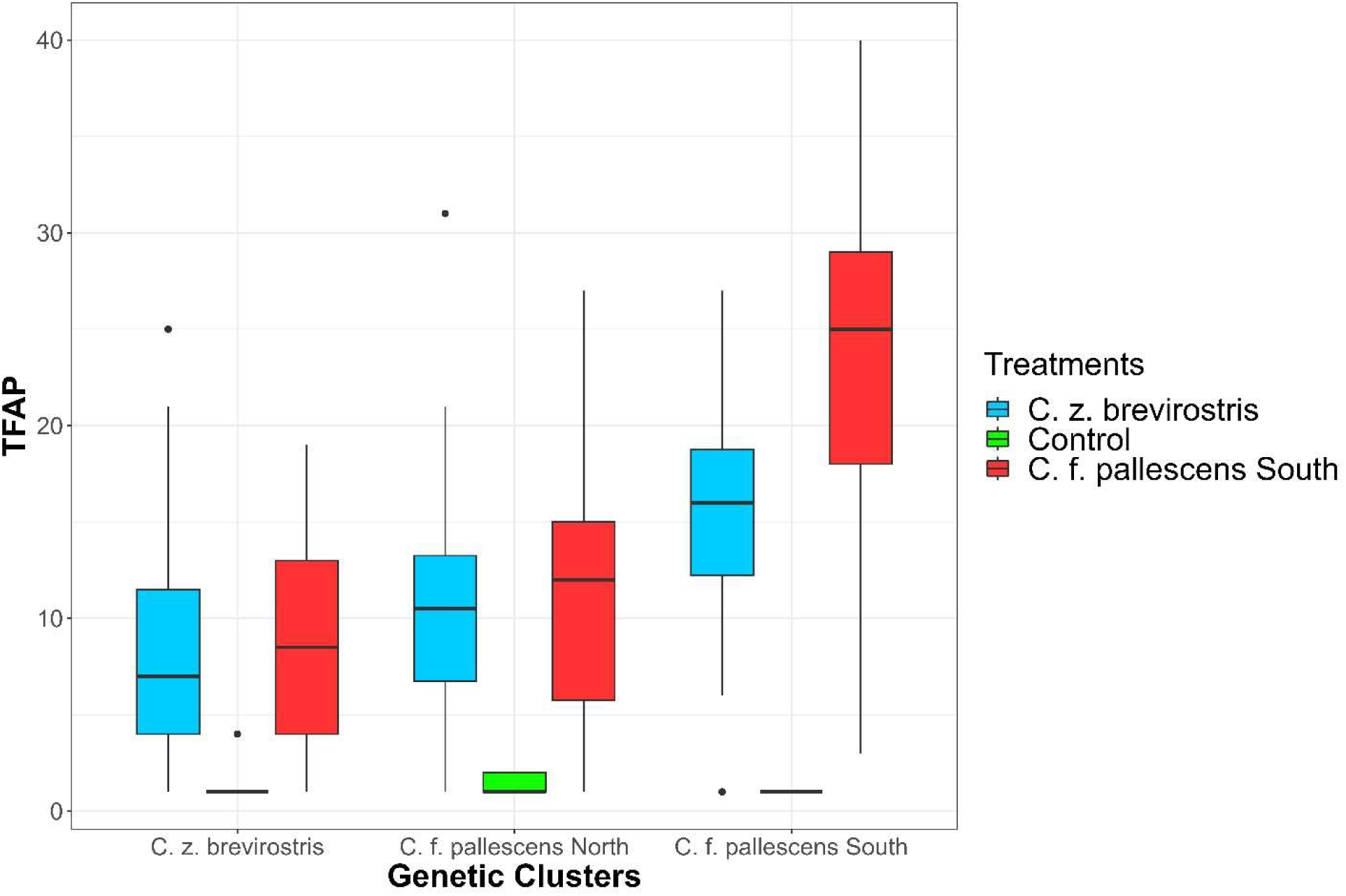
Median and 95% confidence intervals of TFAP per treatment and genetic cluster.

**Figure 6.**
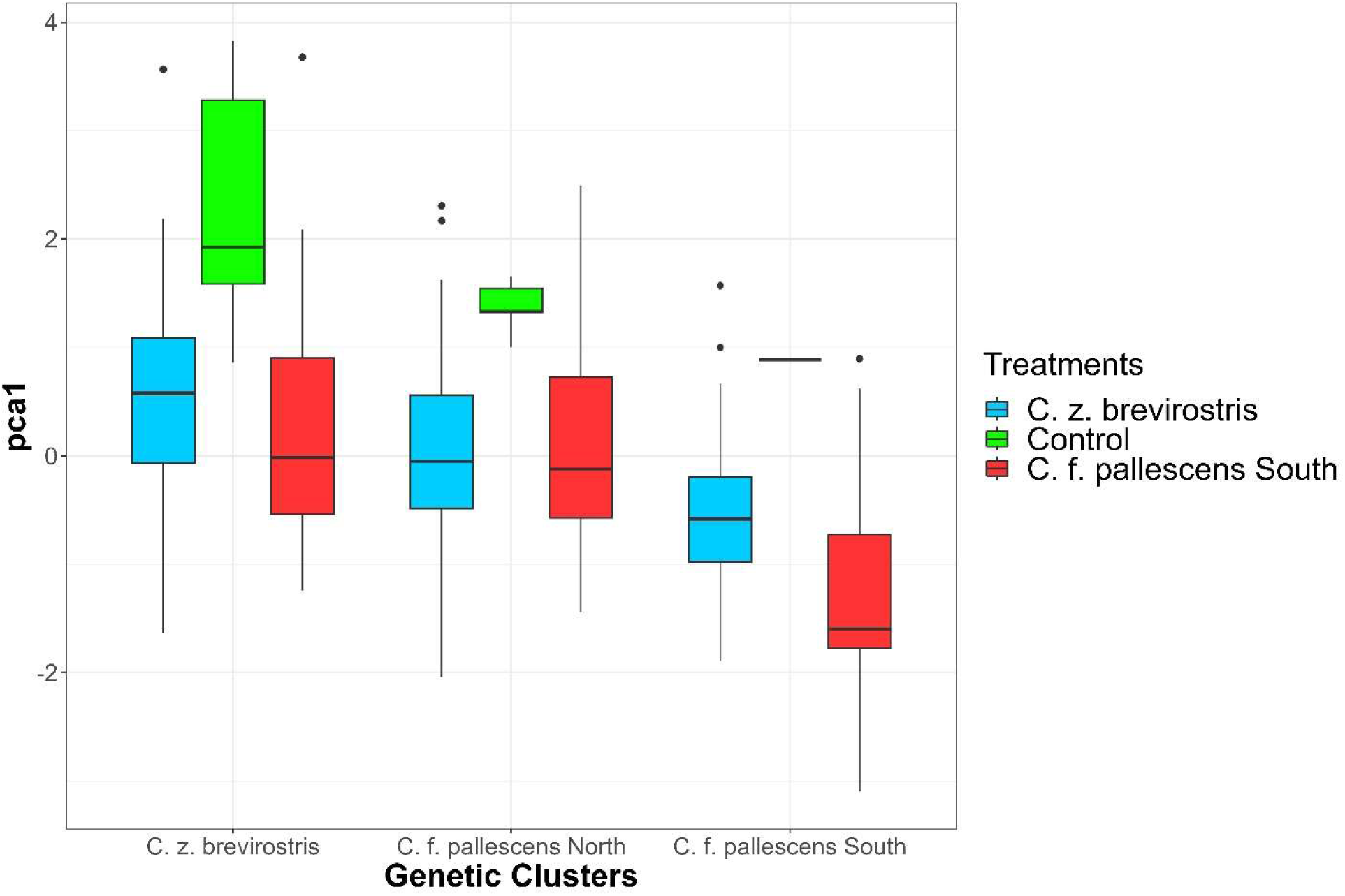
Median and 95% confidence intervals of pca1 per treatment and genetic cluster.

We did observe significant differences between intra- and interspecific treatments in the genetic cluster *C. f. pallescens* South for TFAP (0.27, 95% CI 0.06 to 0.48), supporting our hypothesis H1 (Figures 4-6). However, we did not observe significant differences for pca1 (-0.50, 95% CI -1.20 to 0.21) and minimum distance (-0.61, 95% CI -1.34 to 0.12).

We did not detect significant differences between intra- and interspecific treatments in the genetic cluster *C. z. brevirostris* (Minimum distance=-0.41, 95% CI -0.91 to 0.10; TFAP=0.12, 95% CI -0.04 to 0.29; pca1=-0.21, 95% CI -0.70 to 0.28). However, the negative directionality of the posterior estimates suggests *C. z. brevirostris* exhibited somewhat stronger territorial responses when exposed to the interspecific treatment broadcasting *C. f. pallescens* South songs compared to intraspecific treatments. Although these differences did not reach statistical significance, the pattern was consistent across the dependent variables examined.

### Genetic Clusters and Territorial Aggression

*C. z. brevirostris* showed significantly lower TFAP than the admixed genetic cluster *C. f. pallescens* North (0.17, 95% CI 0.01 to 0.33), supporting our prediction for hypothesis H2. However, we did not observe significant differences for minimum distance (-0.40, 95% CI -0.89 to 0.08), and pca1 (-0.37, 95% CI -0.84 to 0.10).

Contrary to our predictions of hypothesis H2, the genetic cluster *C. f. pallescens* South was significantly closer to the source of stimuli (minimum distance=-1.13, 95% CI -1.66 to - 0.61), registered more TFAP (0.61, 95% CI 0.45 to 0.76), and significantly more territorial aggression (pca1=-1.01, 95% CI -1.51 to -0.50) than *C. f. pallescens* North.

### Predictors of Territorial Aggression

We constructed 28 Bayesian generalized linear models (BGLMs) to identify potential predictors of territorial aggression in *Campylorhynchus* populations in western Ecuador. Gelman-Rubin diagnostic values of 1 for all models and visual inspection of Markov chain Monte Carlo (MCMC) chains indicated satisfactory mixing and convergence (Table 3).

**Table 3.** Summary of variables and model selection diagnostics of BGLMs. Seven models are compared for each response variable. For best models elpd_diff=0. A four-points increment in elpd_diff is considered statistically significant. Percentage of k̂ estimates over 0.7 indicates poor reliability if p_loo is lower than the actual number of model parameters p. GC=Genetic Cluster, AMT=Annual Mean Temperature, AMP=Annual Mean Precipitation.

| Response Variables | Explanatory Variable | Model | elpd_diff | se_diff | elpd_loo | se_elpd_loo | p_loo | se_p_loo | looic | se_looic |
| --- | --- | --- | --- | --- | --- | --- | --- | --- | --- | --- |
| Minimum Distance | All | M7 | 0 | 0 | -553.35 | 22.30846 | 15.75261 | 3.17964 | 1106.699 | 44.61691 |
|  | Treatment*GC | M1 | -4.98212 | 5.073724 | -558.332 | 23.12849 | 11.15425 | 2.881856 | 1116.664 | 46.25698 |
|  | Group Size | M2 | -12.5942 | 8.604696 | -565.944 | 23.77063 | 4.49783 | 1.415562 | 1131.888 | 47.54125 |
|  | Latitude | M3 | -13.9191 | 8.521811 | -567.269 | 25.42755 | 5.899496 | 2.810523 | 1134.538 | 50.85511 |
|  | AMP | M5 | -22.108 | 7.63488 | -575.458 | 24.34925 | 4.420253 | 0.94589 | 1150.915 | 48.6985 |
|  | PS | M6 | -22.5128 | 8.420967 | -575.863 | 25.10408 | 6.283929 | 3.277725 | 1151.725 | 50.20817 |
|  | Null | M0 | -32.8768 | 8.47489 | -586.227 | 22.7305 | 2.460806 | 0.549644 | 1172.453 | 45.46099 |
|  | AMT | M4 | -34.6 | 8.380858 | -587.95 | 23.0853 | 4.965628 | 1.048762 | 1175.899 | 46.1706 |
| Latency | Null | M0 | 0 | 0 | -1094.15 | 9.716572 | 0.461625 | 0.025868 | 2188.306 | 19.43314 |
|  | AMT | M4 | -0.60501 | 0.301076 | -1094.76 | 9.707713 | 0.877938 | 0.05972 | 2189.516 | 19.41543 |
|  | Group Size | M2 | -0.67023 | 0.204009 | -1094.82 | 9.743918 | 0.859344 | 0.06174 | 2189.646 | 19.48784 |
|  | Latitude | M3 | -0.72031 | 0.132835 | -1094.87 | 9.702492 | 0.924106 | 0.063534 | 2189.746 | 19.40498 |
|  | PS | M6 | -0.75246 | 0.119919 | -1094.91 | 9.709884 | 0.966991 | 0.078694 | 2189.811 | 19.41977 |
|  | AMP | M5 | -0.76002 | 0.103988 | -1094.91 | 9.714315 | 0.97845 | 0.084077 | 2189.826 | 19.42863 |
|  | Treatment*GC | M1 | -2.31921 | 1.204773 | -1096.47 | 9.946155 | 2.859622 | 0.18393 | 2192.944 | 19.89231 |
|  | All | All | -5.08032 | 1.386983 | -1099.23 | 10.0241 | 4.961187 | 0.345071 | 2198.466 | 20.0482 |
| TFAP | All | All | 0 | 0 | -796.756 | 33.05152 | 35.84449 | 3.721304 | 1593.512 | 66.10304 |
|  | Treatment*GC | M1 | -21.1294 | 14.59724 | -817.886 | 36.42943 | 20.90954 | 2.240212 | 1635.771 | 72.85886 |
|  | Latitude | M3 | -45.0228 | 19.18995 | -841.779 | 35.80891 | 8.034207 | 0.877175 | 1683.558 | 71.61782 |
|  | Group Size | M2 | -77.7238 | 30.02531 | -874.48 | 38.54313 | 6.93394 | 0.741028 | 1748.96 | 77.08625 |
|  | PS | M6 | -112.113 | 32.42845 | -908.869 | 39.83786 | 9.546634 | 1.220152 | 1817.738 | 79.67572 |
|  | AMP | M5 | -127.735 | 33.59656 | -924.491 | 40.98763 | 9.861306 | 1.282964 | 1848.982 | 81.97527 |
|  | AMT | M4 | -160.564 | 36.86165 | -957.32 | 44.26526 | 10.32227 | 1.4148 | 1914.64 | 88.53051 |
|  | Null | M0 | -175.753 | 41.33556 | -972.51 | 45.99277 | 5.254254 | 0.553503 | 1945.019 | 91.98554 |
| pca1 | All | All | 0 | 0 | -282.01 | 12.33379 | 12.41754 | 1.711707 | 564.0192 | 24.66757 |
|  | Treatment*GC | M1 | -2.69979 | 3.799346 | -284.709 | 11.88801 | 7.210497 | 1.068459 | 569.4188 | 23.77601 |
|  | Latitude | M3 | -5.98548 | 5.228345 | -287.995 | 11.58045 | 3.297005 | 0.577099 | 575.9902 | 23.16091 |
|  | Group Size | M2 | -8.87416 | 6.083844 | -290.884 | 11.60552 | 3.009398 | 0.577818 | 581.7676 | 23.21103 |
|  | PS | M6 | -17.4505 | 6.922087 | -299.46 | 11.35191 | 3.270084 | 0.570916 | 598.9203 | 22.70382 |
|  | AMP | M5 | -19.3207 | 7.049082 | -301.33 | 11.21833 | 3.205714 | 0.543651 | 602.6607 | 22.43665 |
|  | AMT | M4 | -26.6899 | 7.577916 | -308.7 | 11.39647 | 3.22693 | 0.550475 | 617.3991 | 22.79294 |
|  | Null | M0 | -27.3504 | 7.875209 | -309.36 | 11.34949 | 2.220647 | 0.484798 | 618.7201 | 22.69897 |

Leave-one-out cross-validation revealed 27 models with Pareto k values below 0.7, denoting reliability due to consistency when excluding individual samples (Table 3). The minimum distance (M6) model exhibited a higher leave-one-out cross-validation (p_loo) value than the number of predictors (p) and one Pareto k estimate above 0.7, suggesting a distinct subgroup of the *C. f. pallescens* South population maintaining ≥30 m distance during interspecific playback (Table 3).

The full model including all predictors as fixed effects was selected as the best fit for minimum distance, TFAP, and pca1 scores (elpd_diff = 0). The model with treatment x genetic cluster interaction (M1) ranked second-best across response variables (minimum distance elpd_diff = -10.96; total attacks elpd_diff = -29.93; axis 1 scores elpd_diff = -3.68) (Table 3).

Latitude (M3) was the third best and group size (M2) the fourth best predictor of TFAP (elpd_diff = -45.02) and pca1 scores (elpd_diff = -5.98), while the opposite held for minimum distance (elpd_diff = -12.59; -13.91). Precipitation variables (annual mean precipitation M5; precipitation seasonality M6) ranked fifth and sixth across responses, which fit the predictions of hypothesis H3, while annual mean temperature (M4) did not differ from null except for TFAP (elpd_diff = -160.56) (Table 3).

## DISCUSSION

In this study, we aim to address gaps in understanding the factors determining aggressive territorial behavior. To bridge this knowledge gap, we conducted a case study examining intra- and interspecific territorial aggression exhibited by *C. z. brevirostris* and *C. f. pallescens*. Analyses revealed no significant differences between intra- and interspecific treatments for the admixed *C. f. pallescens* North genetic cluster, supporting H1, while *C. f. pallescens* South displayed increased total functional attacks for interspecific treatments, partially supporting H1. No significant differences were detected between treatments for *C. z. brevirostris*. Contrary to predictions in H2, *C. f. pallescens* South showed closer distance, increased total functional attacks, and greater overall territorial aggression (PCA axis 1 scores) compared to *C. f. pallescens* North and *C. z. brevirostris*. Although precipitation models ranked fifth and sixth across response variables, they outperformed null models, lending support for hypothesis H3 that precipitation in particular influences *Campylorhynchus* territorial aggression in Western Ecuador.

### Hybridization and Territorial Aggression

As predicted in our hypothesis H1, the admixed genetic cluster *C. f. pallescens* North exhibited no significant differences between intra- and interspecific treatments, suggesting hybridization influences territorial aggression patterns in this taxa. Our findings support the hypothesis that interspecific territoriality may stabilize the local coexistence of hybridizing species while enabling a faster transition into sympatry than without it (Cowen et al. 2020, Drury et al. 2020). Interspecific territoriality has been strongly associated with hybridization and resource competition during the breeding season (Drury et al. 2020). In birds, hybridization and introgression can promote interspecific territoriality and associate with overlapping breeding resources (Cowen et al. 2020, Drury et al. 2020). Consequently, hybridizing species are more likely to demonstrate interspecific territoriality compared to non-hybrids (Drury et al. 2020). Furthermore, ancestral similarities in territorial signals are maintained and reinforced by selection when interspecific aggression provides an adaptive advantage (Drury et al. 2020). Indices of resource competition positively correlate with interspecific territoriality (Drury et al. 2020).

### Asymmetric Aggression in Campylorhynchus

We detected significantly higher aggression levels in *C. f. pallescens* South compared to *C. z. brevirostris* and *C. f. pallescens* North, evidenced by increased TFAP, shorter average distance to the stimulus source, and overall greater territorial aggression. The asymmetric territorial behaviors displayed by *Campylorhynchus* in western Ecuador are consistent with the asymmetric competition hypothesis (Cowen et al. 2020), with *C. f. pallescens* South as the aggressive dominant genetic cluster, and align with previous research showing larger species tend to display greater dominance and aggression (Miller et al. 2017). However, contrary to initial predictions in hypothesis H2, *C. z. brevirostris* dominance does not appear to drive the higher gene flow from *C. z. brevirostris* toward *C. f. pallescens* North (Montalvo, in prep).

We propose that *Campylorhynchus* asymmetric introgression may stem from sexual selection through female mate choice and male-male interactions (Stein and Uy 2006, Martin and Mendelson 2016). In some cases, the direction of introgression is determined by sex-controlling reproductive decisions. Our results support stronger impacts of asymmetrical reproductive interference over exploitative and agonistic competition (Grether et al. 2017). The relationship between hybridization and territoriality in birds is complex, with ecological, behavioral, and genetic factors involved (Ottenburghs and Slager 2020).

### Group Size and Aggression

We observed a positive correlation between group size and aggression, along with the harshness of the environment — lower annual mean precipitation and higher precipitation seasonality. This suggests that in Western Ecuador, *C. fasciatus* may exhibit an increase in both group size and aggression under conditions of reduced resource availability.

Studies have shown that group size can affect the level of aggression and territoriality in cooperative breeding birds (Lukas and Clutton-Brock 2012). For example, in some species, larger groups may be more aggressive towards intruders and defend larger territories than smaller groups (Karell et al. 2011, Lukas and Clutton-Brock 2012). However, many studies of cooperatively breeding birds have found no effect of group size on reproductive success, contrary to predictions of most adaptive hypotheses (Magrath 2001). Group size has a reduced effect on success when conditions for breeding are good, such as in good environmental conditions or groups with older breeders (Magrath 2001).

A recent study showed opposing ecogeographic patterns of aggression, and social group size along a steep environmental gradient in two congeneric cooperatively breeding species of fairywrens (Maluridae) (Johnson et al. 2023). *Malurus assimilis* (Purple-backed fairywrens), which have helpers that increase group productivity, have larger groups in hot and dry environments and smaller groups in cool and wet environments (Johnson et al. 2023). However, *Malurus cyaneus* (Superb Fairywrens), which have helpers that do not enhance group productivity despite the presence of alloparental care, showed the opposite trend (Johnson et al. 2023). Thus, it is suggested that differences in the costs and benefits of sociality may contribute to these opposing eco-geographical patterns (Johnson et al. 2023). The observed patterns of group size among *C. fasciatus* across the precipitation gradient within western Ecuador align with those previously recorded for *Malurus assimilis*, whereby larger group sizes are found in drier areas.

### Environment and Aggression

The mechanisms through which climate could affect territorial aggression are multiple. Environmental factors such as climate change might affect aggression and territoriality in birds (DeMoranville et al. 2019, 2020). In our study, precipitation was the climatic variable most relevant to explain the aggression of *C. z. brevirostris* and *C. fasiatus*. To date, there has been a lack of research on how intra and interspecific aggression may be influenced by rainfall. However, there is evidence to suggest that it may affect aggressive territorial behavior through resource availability or habitat quality (Zack and Stutchbury 1992, Cain et al. 2015, França et al. 2020). As resources become scarce, it is conceivable that animals may become territorial and engage in aggressive behavior to protect their territory and ensure they have adequate access to resources (Zack and Stutchbury 1992, Cain et al. 2015, França et al. 2020). To our knowledge, this is the first study linking annual mean precipitation as a significant factor explaining aggressive territoriality in birds.

Despite the lack of significant differences between the model including annual mean temperature and the null model for any of the response variables, we cannot dismiss its potential effect on aggressive territoriality. It is worth noting that past studies have primarily explored the impact of temperature on aggressive behaviors. Temperature affects food availability and consequently, energy expenditures and coping behavioral adaptations (Soma 2006). Temperature could also affect the nature of social interactions by inducing changes in the neuroendocrinological mechanisms that underpin key social traits. In some cases, such mechanisms are already fine-tuned to respond to extrinsic cues (Moss and While 2021). These effects start to take effect during the early stages of group formation when minute temperature-induced behavioral changes (Anderson et al. 2000) (such as length of residency, thermoregulatory aggregation, territorial aggression, and natal dispersal) modulate the frequency and duration of associations between kin and non-kin (Moss and While 2021).

While the BGLM models presented in our study exhibit substantial effects, it is important to acknowledge the possibility that the observed effects could be attributed, to some extent, to the size of our sample. Even in the case of well-specified models and models with different predictions, small data (say less than 100 observations) makes estimating uncertainty in cross-validation less reliable (Lee and Song 2004, Sivula et al. 2022). Penalization and shrinkage methods produced unreliable clinical prediction models, especially when the sample size is small (Riley et al. 2021). When the differences between models are small, or models are slightly misspecified, even larger data sets are needed for well-calibrated model comparison (Lee and Song 2004, Sivula et al. 2022). Annual mean temperature yielded negligible and incongruous outcomes across the response variables, which could be attributed to the small sample size, and thus requires further examination. Further research should be conducted with a larger sample to validate the results and minimize the potential influence of sample size on the observed effects.

### Potential Mechanisms Determining Aggression

The variability of aggressive behaviors, both within and between individuals, can present challenges when attempting to identify patterns of determinant factors that influence this behavior. For instance, *Thryomanes bewickii* (Bewick’s wrens) and *Troglodytes aedon* (House Wrens) defend nonoverlapping territories at many locations, but one study found extensive territory overlap and little interspecific aggression (Kroodsma 1973), which suggests that aggressive territoriality is facultative in this species pair. The task of identifying behavioral patterns in birds becomes increasingly challenging in the presence of significant variability within and among subjects (Samia et al. 2017). The complex and multifactorial nature of such variability presents challenges in the identification of patterns, as it can obscure relevant differences between groups or individuals. Genetic factors can also play a role in aggression and territoriality in birds, as studies have shown that basal metabolic rate (BMR) is heritable in birds and can be modulated by genetic factors (Tieleman et al. 2009).

Territorial aggression in birds can vary considerably across the year in the northern hemisphere (Landys et al. 2007, Merritt et al. 2018). Conversely, year-round territorial behavior is more prevalent among tropical birds (Hau et al. 2000, Fedy and Stutchbury 2005). This distinction is an important consideration for the understanding of avian behavior in different ecozones. We suspect that *Campylorhynchus* in western Ecuador keep territories around the year. Communal signaling such as duets — as in *Campylorhynchus* — has been associated with year-around territoriality and stable social bonds (Tobias et al. 2016). During our fieldwork for the study, we were able to observe active nests, breeding patches, and juveniles for the study species. These observations support the concept of year-round breeding for these species.

## CONCLUSION

We investigated intra- and interspecific aggressive territorial behaviors exhibited by *C. z. brevirostris* and *C. f. pallescens*, as well as potential associations with spatial climate variability in western Ecuador. Our findings support the hypothesis that hybridizing species tend to stabilize local coexistence through territorial aggression, as evidenced by comparable territorial aggression responses to intra- and interspecific stimuli in the admixed *C. f. pallescens* North genetic cluster.

We showed that the level of aggression in *Campylorhynchus* varied significantly between the different subspecies. Specifically, *C. f. pallescens* South exhibited significantly higher aggression levels than both *C. z. brevirostris* and *C. f. pallescens* North. Contrary to our initial hypothesis, our study found that the dominance of *C. z. brevirostris* was not the driving force behind the observed introgression patterns between the two species. Therefore, this finding warrants further investigation to determine the underlying mechanisms behind introgression. We suggest that seasonal constraints on habitat quality exert selective pressures for increased territorial aggression as a mechanism to ensure adequate resource availability.

The results suggest that multiple predictors contribute to aggressive territorial behavior in birds, including group size, genetic clusters, latitude, and precipitation. The study also found that precipitation was the climate factor more important to explain the aggression of *C. z. brevirostris* and *C. f. pallescens*. The complex and multifactorial nature of territorial aggression in birds and the variability of aggressive behaviors within and among individuals suggest the need for further study into the underlying mechanisms behind introgression and asymmetrical responses of aggression. Further research in this area will be important to uncover the specific mechanisms behind introgression, with a particular focus on the relationship between hybridization, territoriality, and environmental factors.

## Notes

### Competing Interest Statement

The authors have declared no competing interest.

